# HlyF/CprA modulate bacterial extracellular vesicle properties and mediate efficient host cell cargo delivery

**DOI:** 10.1101/2025.03.27.645671

**Authors:** Frédéric Taieb, Laure David, Annie Frelet-Barrand, Camille Pin, Audrey Goman, Asja Garling, Christelle Marrauld, Marine Colson, Marie Pénary, Eric Oswald

**Author notes:** These authors contributed equally (Frédéric Taieb, Laure David).

## Abstract

HlyF, a cytoplasmic short-chain dehydrogenase/reductase encoded by ColV-like virulence plasmids in *Escherichia coli*, and its ortholog CprA, encoded on the chromosome of *Pseudomonas aeruginosa*, drive the production of a distinct population of bacterial outer membrane vesicles (OMVs). These vesicles were previously shown to disrupt autophagic flux and amplify host inflammatory responses, underscoring their functional divergence from canonical OMVs. Here, we demonstrate that HlyF/CprA-dependent OMVs also display enhanced cargo delivery efficiency through a dynamin-independent internalization pathway involving direct fusion with the host cell plasma membrane. This non-canonical uptake mechanism differentiates these vesicles from conventional bacterial OMVs and promotes efficient cytosolic delivery of vesicular cargo. Collectively, our findings reveal a previously unrecognized role for the HlyF/CprA enzyme family in modulating OMVs biogenesis and functional properties, and identify a novel OMV-mediated mechanism contributing to bacterial virulence.

## Introduction

Gram-negative bacteria, like all cells, release extracellular vesicles (EVs). Among the different EV subtypes, vesicles that bud from the outer membrane are termed outer membrane vesicles (OMVs). These nanoscale particles serve as dedicated transport vehicles for a diverse repertoire of bacterial components derived from both the outer membrane and the periplasm. OMVs and their cargo include lipids, lipopolysaccharides (LPS), outer membrane proteins (OMPs), and periplasmic constituents, notably virulence-associated factors such as toxins (Schwechheimer and Kuehn 2015; Jan 2017; Juodeikis and Carding 2022; Bielaszewska et al. 2017; Kunsmann et al. 2015; Murase et al. 2016). By displaying surface-exposed pro-inflammatory antigens and microbial/pathogen-associated molecular patterns (M/PAMPs), OMVs also potently modulate innate and adaptive immune responses and can elicit severe inflammatory outcomes, including sepsis (Park et al. 2010; Santos et al. 2018; Eren et al. 2020; Vanaja et al. 2016). Importantly, numerous studies have demonstrated that eukaryotic cells internalize bacterial vesicles through endocytic pathways, allowing OMVs to efficiently deliver not only membrane-associated components (lipids, OMPs, PAMPs, and LPS) but also toxins and other periplasmic cargo directly into host cells, thereby positioning OMVs as a central mechanism of toxin delivery during host–pathogen interactions (Pathirana and Kaparakis-Liaskos 2016; O’Donoghue and Krachler 2016; Sabatke et al. 2024; Bielaszewska et al. 2017). Accordingly, OMVs are now recognized as key mediators of bacterial communication and virulence.

HlyF and CprA are cytoplasmic enzymes and are excluded from OMVs. Nevertheless, they drive the production of a specific subset of OMVs that are capable of blocking autophagic flux in intoxicated eukaryotic cells (David et al. 2022; Goman et al. 2025; Murase et al. 2016). Because autophagy contributes to negative feedback regulation of inflammasome activation, OMV-dependent inhibition of autophagy exacerbates the non-canonical inflammatory response triggered by OMV-mediated PAMPs delivery, ultimately leading to host cell death (Lapaquette et al. 2015; David et al. 2022; Santos et al. 2018; Pin, David, and Oswald 2023; Goman et al. 2025). We have recently shown that HlyF and CprA belong to a family of shorty chain dehydrogenase/reductase (SDR) enzymes that modify lipids. Through this enzymatic activity, HlyF and CprA generate outer membrane toxic lipid species that are responsible for the deleterious biological effects of the resulting OMVs (Goman et al. 2025). Consistently, expression of HlyF and CprA has been linked to bacterial pathogenicity in *Escherichia coli* and *Pseudomonas aeruginosa* in both human and animal (Chagneau et al. 2023; Goman et al. 2025; Murase et al. 2016). In this study, we demonstrate that OMVs produced by HlyF/CprA-expressing bacteria display enhanced cargo delivery efficiency into host cells. Mechanistically, HlyF/CprA promotes OMV entry through a dynamin-independent pathway involving direct fusion with the host cell plasma membrane, thereby enabling efficient delivery of vesicular virulence factors and reinforcing the central role of OMVs in bacterial pathogenicity.

## Results

### HlyF expression does not alter OMV composition, size distribution, or TEM-1 cargo loading

TEM-1 ß-lactamase serves as an ideal reporter for tracking OMV uptake because its enzymatic activity can be readily quantified in eukaryotic cells, providing a direct measure of cargo delivery. In Gram-negative bacteria, TEM-1 β-lactamase is naturally translocated into the periplasmic space via the Sec-system pathway, where it acquires its active form following cleavage of the N-terminal signal peptide and oxidative folding (Kaderabkova et al. 2022). Consequently, OMVs, which are filled with periplasmic contents, naturally incorporate TEM-1 β-lactamase as cargo (Figure 1A). We purified OMVs loaded with TEM-1 ß-lactamase from *E. coli* BL21(DE3) strain co-transformed with plasmids encoding either wild-type HlyF (HlyF^wt^) or an inactive HlyF mutated in the SDR catalytic site (HlyF^mut^), together with the high-copy pUC19 plasmid expressing TEM-1 β-lactamase. Western blot quantification of TEM-1 β-lactamase in both OMV types, using the outer membrane protein A (OmpA) as a loading control, revealed comparable protein levels, indicating that HlyF does not affect vesicle’s enzyme incorporation (Figure 1B). Enzymatic activity was then determined by measuring the absorbance of hydrolyzed chromogenic substrate nitrocefin over time in presence of purified OMVs. Both the reaction slope and the total enzymatic activity showed that OMVs from *E. coli* expressing either wild-type or mutant HlyF, which contain equivalent amounts of TEM-1 β-lactamase as confirmed by western blot analysis, exhibited comparable enzyme activity (Figure 1C and Figure S1A). Microscopic observation revealed a consistent similar vesicle structure, as expected for OMV formation (Figure S1B). In addition, Coomassie blue and silver staining of SDS-PAGE gels shown comparable protein profiles between HlyF^wt^ and HlyF^mut^ OMVs (Figure S1C).

**Figure 1.**
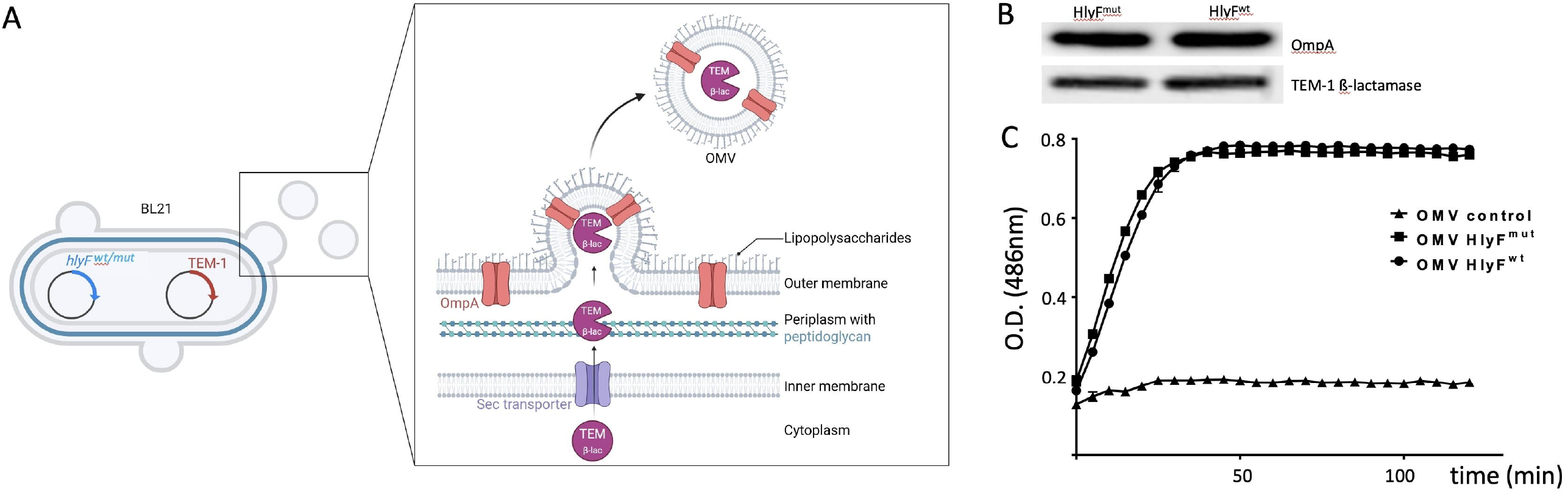
Method of measurement of TEM-1 β -lactamase in host cells. (A) Schematic representation of OMV loaded with TEM-1 ß-lactamase: *E. coli* BL21(DE3) co-transformed with plasmids encoding TEM-1 β-lactamase and either wild-type (*WT*) or mutant (*mut*) *hlyF* produce OMVs containing TEM-1 β-lactamase (magenta) and OmpA (pink barrel). Created with BioRender.com; (B) Western blot detection of OmpA and TEM-1 β-lactamase from 0.5µg of purified OMVs from BL21 strains expressing wild type (HlyF^wt^) or inactive mutated (HlyF^mut^) HlyF; (C) Representative kinetic assay of nitrocefin hydrolysis measured by O.D. at 486nm over time upon incubation with indicated OMVs (OMVs control correspond to OMVs produced in absence of pUC19 encoding TEM-1 ß-lactamase) at the same concentration (see also Figure S1A)

OMV size distribution and concentration were then analyzed by multiangle dynamic light scattering (MADLS). This state-of-the-art technique, in contrast to the single detection angle of conventional dynamic light scattering (DLS), combines measurements acquired at multiple scattering angles enabling estimation of particle concentration in addition to hydrodynamic diameter with robust and accurate determination of particle size from 0.3 nm to 15 µm (Naiim et al. 2015; Normak et al. 2023). Moreover, MADLS allows precise measurement of OMV concentration whereas nanoparticle tracking analysis (NTA) fails to detect vesicle under 60-70 nm (Bachurski et al. 2019; Arab et al. 2021). These advantages are particularly relevant for small extracellular vesicles such as OMVs which start at 20 nm in size (Schwechheimer and Kuehn 2015; Juodeikis and Carding 2022) and in our study, measured 25-30 nm in diameter. MADLS measurements confirmed the size distribution previously reported by David *et al*. (2022) and further allowed us to determine OMV particle concentrations in our preparations. Size distributions were comparable between HlyF^wt^ and HlyF^mut^ OMVs, with mean diameters of 29.6 ± 2.6 nm and 24.8 ± 2.2 nm, respectively, showing no statistically significant difference between the two populations (Welch’s t-test, p = 0.074). Across all samples, the mean diameter was of 28.2 ± 1.3 nm (standard error of the mean, n=8). A second, larger OMVs population with diameters >50 nm was also detected (Figure 2A); however, this population accounted for only a minor fraction (0.27%) of the total OMVs populations (Figure S2A and S2B). At a normalized protein concentration of 1mg/mL, the concentration of OMVs with diameters <50 nm was estimated at 2.7 × 10^15^ ± 3.5 × 10^14^ vesicles/mL for HlyF^wt^ OMVs and 3.0 × 10^15^ ± 2.2 × 10^15^ vesicles/mL for HlyF^mut^ OMVs, with no statistically significant difference between the two types of OMV (Welch’s t-test, p = 0.81) (Figure 2B and Figure S2A and S2B). Overall, these results demonstrate comparable particle-to-proteins ratios between HlyF^wt^ and HlyF^mut^ OMVs samples, with an average concentration of 2.85 × 10^15^ OMVs per mg of protein (± 4.3 × 10^14^, standard error of the mean, n= 8).

**Figure 2.**
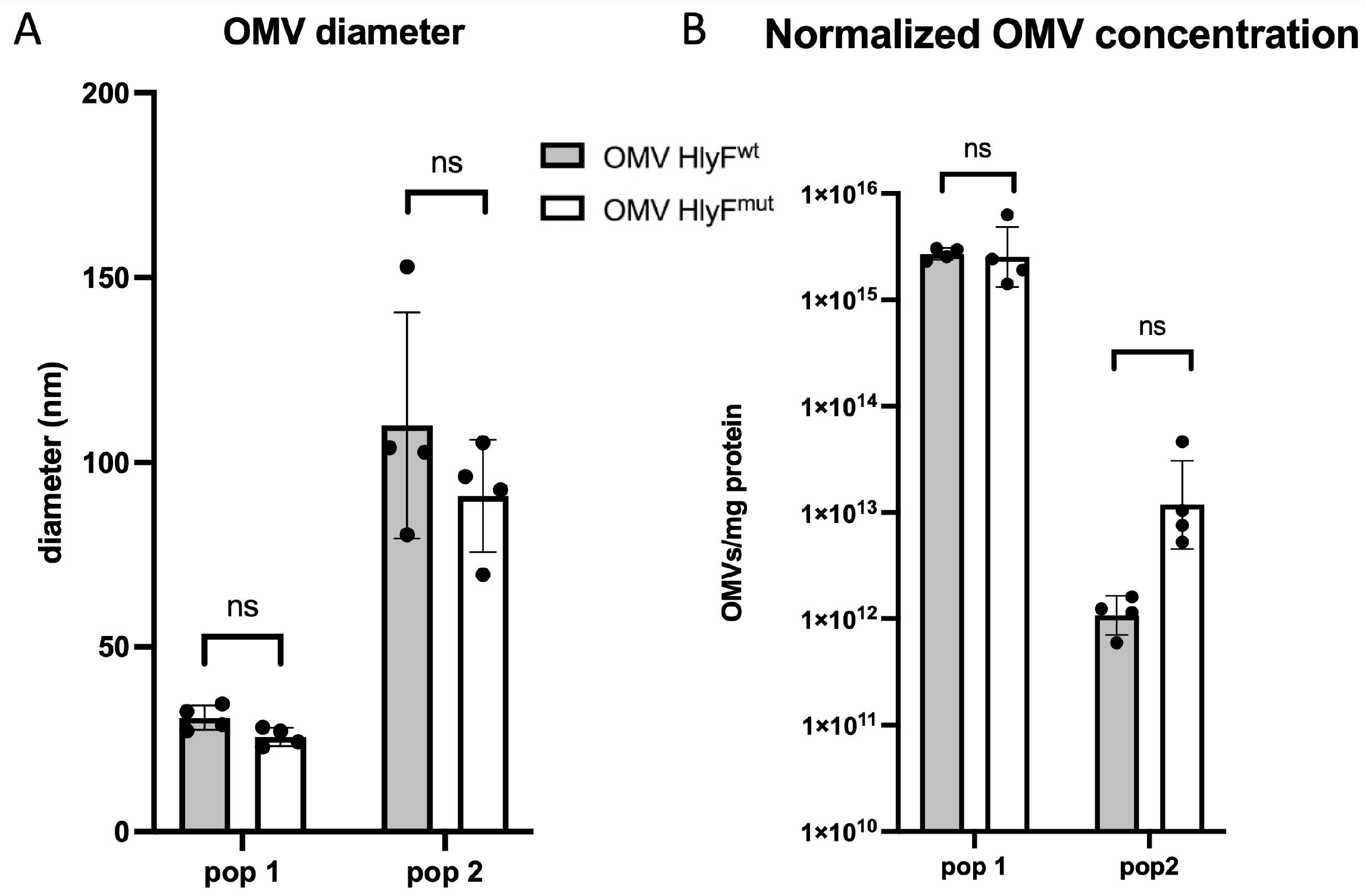
Multi-angle dynamic light scattering (MADLS) analysis. HlyF^wt^ (grey bar) and HlyF^mut^ (white bar) OMVs (A) diameter and (B) normalized concentration (OMV to protein ratio); pop 1 and pop 2 correspond to 2 populations of OMVs <50 nm (pop 1) and >50 nm (pop2). Data are represented as mean ± standard deviation (SD), n = 4 independent experiments; ns: non significative difference, Welch’s t-test (see also Figure S2A)

To further assess whether the ultracentrifugation-based purification procedure introduced any bias in OMV size distribution or concentration, we compared OMV preparations obtained after ultracentrifugation (uc) with those subjected to an additional OptiPrep density gradient purification step (opti). The fractions displayed highly similar size distributions (Z-average diameters of 34.05 nm (uc) vs 32.45 nm (opti) for HlyF^wt^ OMVs and 27.15 (uc) vs 28.20 nm (opti) for HlyF^mut^ OMVs) and concentrations (4.82 × 10^15^ (uc) vs 5.63 × 10^15^ (opti) for and 4.35 × 10^15^ vs 2.26 × 10^15^ for HlyF^wt^ and HlyF^mut^ OMVs respectively) (Figure S2C), confirming that ultracentrifugation alone provides representative and sufficiently pure OMV preparations for subsequent experiments.

Overall, these data establish that OMVs produced by HlyF^wt^ and HlyF^mut^ -expressing bacteria constitute highly comparable vesicle populations with respect to biochemical composition and biophysical properties, providing a robust experimental framework for directly comparing vesicles uptake and cargo delivery into host cells.

### HlyF enhances OMV cargo delivery kinetics in mammalian cells

To investigate the kinetics of cargo delivery into eukaryotic cells, we used a reporter system based on detection of TEM-1 β-lactamase activity, using the fluorescence resonance energy transfer (FRET) substrate CCF4-AM (LiveBLAzer™). This substrate consists of a cephalosporin core linking 7-hydroxycoumarin to fluorescein. Enzymatic cleavage of the ß-lactam ring of cephalosporin disrupts CCF4 FRET, resulting in a fluorescence shift from green to blue enabling quantification of TEM-1 β-lactamase cargo delivery by OMVs (Figure 3A). Purified OMVs containing a normalized quantity of TEM-1 β-lactamase were added to HeLa cells preloaded with CCF4. The ratio of blue to green fluorescence was monitored over time using a microplate spectrophotometer, allowing precise quantification of TEM-1 β-lactamase delivery into host cells (Figure 3B). Total enzyme delivery was assessed by calculating the area under the curve (AUC). AUC analysis revealed that OMVs derived from the *E. coli* BL21(DE3) strain efficiently delivered TEM-1 β-lactamase into HeLa cells. Notably, OMVs produced by bacteria expressing wild-type HlyF (HlyF^wt^) resulted in significantly greater accumulation of TEM-1 β-lactamase compared with OMVs from the catalytic mutant strain (HlyF^mut^) (Figure 3C). Moreover, the steeper slope of the blue/green fluorescence ratio for HlyF^wt^ OMVs indicates an accelerated rate of enzyme accumulation, accounting for their enhanced overall delivery efficiency relative to HlyF^mut^ OMVs (Figure 3D). We next examined whether this enhanced cargo delivery was conserved in OMVs produced by pathogenic bacteria that endogenously express HlyF^mut^ or its ortholog CprA. These included the uropathogenic *E. coli* clinical isolate ECC166 and its isogenic hlyF deletion mutant [12] as well as the *P. aeruginosa* PAK strain, which encodes *cprA*, and its corresponding isogenic *cprA* mutant [9]. Consistent with results obtained using the BL21(DE3) model strain, AUC analysis revealed a HlyF/CprA-dependent increase in CCF4 cleavage by OMVs from both pathogenic species (Figure S3). Because all OMVs preparations were normalized for equivalent TEM-1 β-lactamase content, these data demonstrate that OMVs derived from HlyF/CprA-expressing bacteria promote enhanced intracellular cargo accumulation. To determine whether this phenomenon extends beyond transformed epithelial cell lines, we next analyzed primary mouse bone marrow-derived macrophages (BMDMs). Enhanced delivery of TEM-1 β-lactamase by HlyF^wt^ OMVs was similarly observed in CCF4-loaded BMDMs (Figure 3E), validating this effect in a physiologically relevant immune cell context. Finally, to exclude any contribution from free or surface-exposed TEM-1 β-lactamase, experiments were repeated using pronase-treated OMVs. The efficiency of pronase digestion was confirmed by protein profiling and western blot analysis of OmpA and TEM-1 β-lactamase, showing that both the transmembrane proteins and luminal cargo remained protected from proteolytic cleavage (Figure S4A and S4B) whereas other proteins, mainly those of high molecular weight were efficiently digested. In this CCF4 assay, we used 2.5-fold more HlyF^mut^ OMVs than HlyF^wt^ OMVs (25µg/mL and 10µg/mL respectively) to compensate for the lower ß-lactamase activity observed in cells treated with HlyF^mut^ OMVs. While a slight decrease of AUC is observed in protease-digested OMVs, comparable AUC values were obtained under these conditions, indicating that CCF4 cleavage is primarily driven by OMV-associated TEM-1 β-lactamase cargo (Figure S4C).

**Figure 3.**
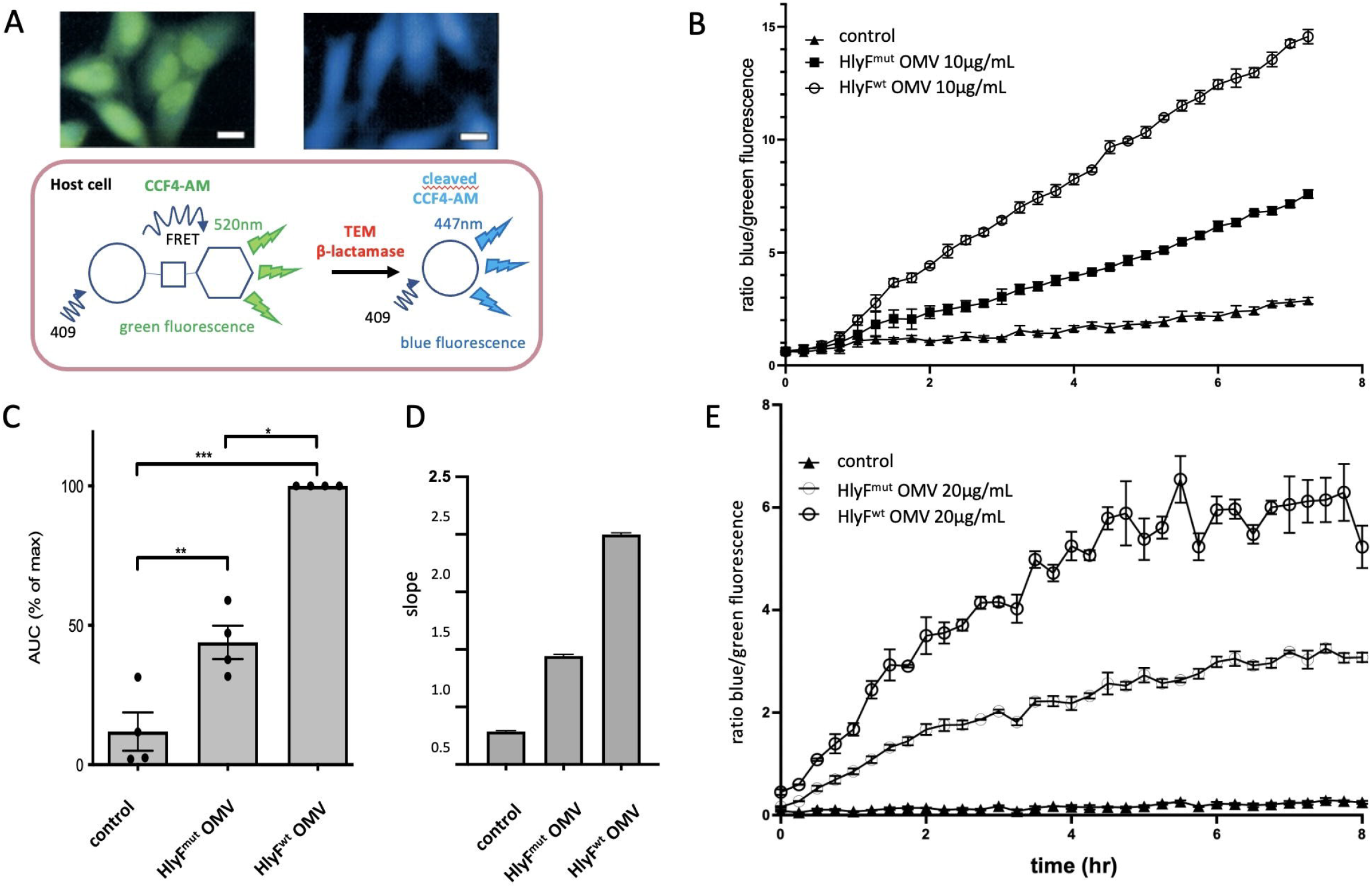
HlyF promotes accumulation of TEM-1 TEM-1 β-lactamase in eukaryotic cells. (A) Principle of CCF4-AM assay: intact CCF4 emits green fluorescence, whereas cleavage by TEM-1 β-lactamase delivered through internalized OMVs results in blue fluorescence due to FRET loss; (B) Time course of blue/green fluorescence ratios in HeLa cells treated with 10µg/mL of OMVs from BL21(DE3) expressing wild type (HlyF^wt^) or mutated HlyF (HlyF^mut^). Data are representative of technical triplicate experiments and are represented as mean ± SD; (C) Quantification of intracellular TEM-1 β-lactamase activity expressed as the area under the curve (AUC) from panel A, calculated from four independent experiments with 10µg/mL OMVs. Data are represented as mean ± standard error of the mean (sem); statistical significance assessed by one-way ANOVA with Bonferroni correction (*p < 0.05, **p < 0.01, ***p < 0.001). (D) Slopes of blue/green fluorescence curves from panel B determined by linear regression. (E) Time course of blue/green fluorescence ratio in mouse bone marrow-derived macrophages treated with 20µg/mL of OMVs from BL21(DE3) expressing wild type (HlyF^wt^) or mutated HlyF (HlyF^mut^). Data are representative of technical triplicate experiments and are represented as mean ± SD.

### Enhanced cargo delivery by HlyF OMVs is independent of autophagy inhibition

Given that HlyF^wt^ OMVs have been shown to block autophagosome-lysosome fusion (Pin, David, and Oswald 2023; David et al. 2022), we hypothesized that degradation of these OMVs by autophagy might be disrupted. Such inhibition could stabilize OMV cargo, including TEM-1 β-lactamase, thereby increasing CCF4-AM cleavage and the associated AUC values. To test this hypothesis, we treated cells with the autophagy inhibitors chloroquine and bafilomycin (Stamenkovic et al. 2019). If the increased AUC observed with HlyF^wt^ OMVs resulted from TEM-1 stabilization due to autophagic blockage (David et al. 2022), then autophagy inhibitors would be expected to increase AUC in HlyF^mut^ OMV-treated cells to levels comparable to those observed with HlyF^wt^ OMVs. However, addition of these pharmacological inhibitors did not increase AUC values in HlyF^mut^ OMV-treated cells (Figure 4A and 4B), indicating that the enhanced TEM-1 β-lactamase activity is not attributable to autophagic dysfunction but rather reflects improved cargo delivery by OMVs produced by bacteria expressing the catalytically active HlyF.

**Figure 4.**
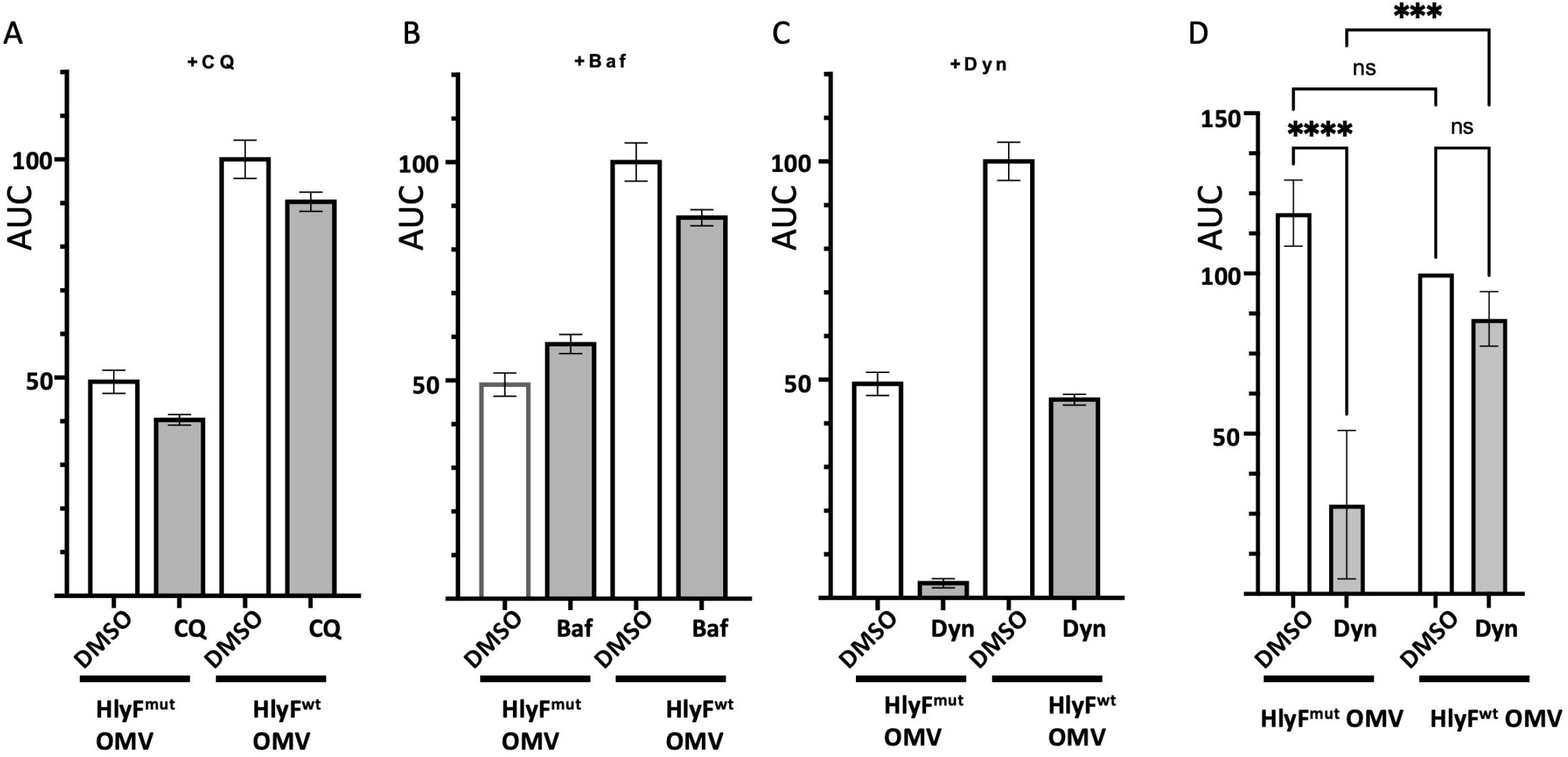
Pharmacological inhibition of OMVs produced by BL21(DE3) expressing wild-type or inactive HlyF. HeLa cells were treated with 10µg/mL OMVs and intracellular TEM-1 β-lactamase activity was quantified as AUC. (A-C) AUC values for cells incubated with HlyF^wt^ or HlyF^mut^ OMVs in the presence of DMSO (white bars) or pharmacological inhibitors (grey bars): (A) chloroquine 100µM (CQ), (B) bafilomycin 100nM (Baf), and (C) Dynasore 80µM (Dyn); AUC are represented as the mean and SD from technical triplicate; (D) AUC values were calculated from three independent experiments using HlyF^wt^ OMVs (2µg/mL) or HlyF^mut^ OMVs (5µg/mL) in the presence of Dynasore or DMSO. Data are expressed as percentage relative to the HlyF^wt^ + DMSO condition. AUC are represented as the mean ± SEM. Statistical analysis by one-way ANOVA with Bonferroni correction; ns, not significant; ***p < 0.01; ****p < 0.001.

### HlyF enables dynamin-independent OMV entry into host cells

OMVs derived from *E. coli* have been reported to enter host cells through both clathrin- and caveolin-dependent endocytosis pathways (Bielaszewska et al. 2017; Sabatke et al. 2023; Canas et al. 2016; Krsek et al. 2023; Thapa et al. 2023; Kesty et al. 2004). Formation of endocytic vesicles requires coordinated activity of multiple host proteins that promote plasma membrane invagination and vesicle scission, a process mediated by the GTPase dynamin, which catalyzes membrane fission to generate early endosomes (Doherty and McMahon 2009; Kumari, Mg, and Mayor 2010). To investigate the HlyF-dependent mechanism of OMV internalization, we used Dynasore, a dynamin GTPase inhibitor that blocks both clathrin- and caveolin-mediated endocytosis (Macia et al. 2006; Marks et al. 2001). CCF4-loaded cells were treated or not with Dynasore and exposed to either HlyF^wt^ or HlyF^mut^ OMVs containing equivalent amounts of TEM-1 cargo. Consistent with previous observations for OMVs produced by *E. coli strains lacking hlyF* (Bielaszewska et al. 2017; Sabatke et al. 2023; Canas et al. 2016; Krsek et al. 2023; Kesty et al. 2004), Dynasore inhibited the entry of OMVs from bacteria expressing the inactive HlyF variant. In contrast, although Dynasore treatment reduced the AUC of HlyF^wt^ OMVs relative to untreated controls, substantial internalization persisted in inhibitor-treated cells (Figure 4C). The residual TEM-1 β-lactamase activity detected in the presence of Dynasore could reflect non-specific intrinsically higher uptake efficiency of HlyF^wt^ OMVs. To exclude this possibility, we repeated the assay using 2.5-fold fewer HlyF^wt^ OMVs, adjusted to yield comparable AUC values between HlyF^wt^ and HlyF^mut^ OMVs in untreated cells. Under these normalized conditions, Dynasore inhibited HlyF^wt^ OMV entry by nearly 80%, whereas internalization of HlyF^wt^ OMVs was reduced by only ∼10% (Figure 4D). Together, these data indicate that HlyF promotes OMV entry into host cells via an alternative, dynamin-independent internalization pathway. Notably, the observation that higher concentrations of HlyF^wt^ OMVs exhibited partial inhibition (∼50%, Figure 4C), whereas lower concentrations were minimally affected (∼10%, Figure 4D), suggests that at elevated OMV levels, uptake may partially rely on dynamin-dependent endocytosis once the HlyF-dependent entry pathway becomes saturated.

### HlyF^wt^ OMVs enter cells through direct plasma membrane fusion

To investigate the mechanism of HlyF^wt^ OMVs internalization into eukaryotic cells, we labeled OMVs membranes with the lipophilic dye DiI, which spontaneously integrates into lipid bilayers. After confirming that HlyF^mut^ and HlyF^wt^ OMVs incorporated DiI with comparable efficiency (Figure S5), HeLa cells were exposed to the labeled vesicles. Cells treated with HlyF^mut^ OMVs exhibited punctate intracellular fluorescence, indicating that the dye remained confined within intact internalized vesicles, consistent with uptake via endocytic compartments. In contrast, cells exposed to HlyF^wt^ OMVs exhibited faint and diffuse fluorescence that was nearly indistinguishable from background, a pattern consistent with redistribution of the lipophilic dye into host cell membranes following vesicle–plasma membrane fusion. Quantitative numeration of dot puncta confirmed the distinct staining patterns observed in cells treated with the two types of OMVs (Figure 5A). To further test this hypothesis, we examined the involvement of plasma membrane lipid raft microdomains, which have been shown to mediate *P. aeruginosa* OMV fusion (Bomberger et al. 2009). We used FITC-labeled cholera toxin B subunit (CtxB-FITC), a well-established lipid raft marker (Harder et al. 1998). In untreated cells and in cells exposed to HlyF^mut^ OMVs, CtxB-FITC fluorescence appeared as discrete clusters, visible as green puncta corresponding to organized lipid rafts. In contrast, cells treated with HlyF^wt^ OMVs exhibited fewer and more diffusely distributed CtxB-FITC clusters, indicative of altered raft organization (Figure 5B). Because dispersion of CtxB-FITC foci reflects changes in plasma membrane lipid dynamics (Harder et al. 1998), these observations support a model in which, unlike HlyF^mut^ OMVs that enter cells via endocytosis without perturbing membrane organization, HlyF^wt^ OMVs directly fuse with the plasma membrane, inducing lipid reorganization and enabling efficient cargo delivery directly in the cytosol.

**Figure 5.**
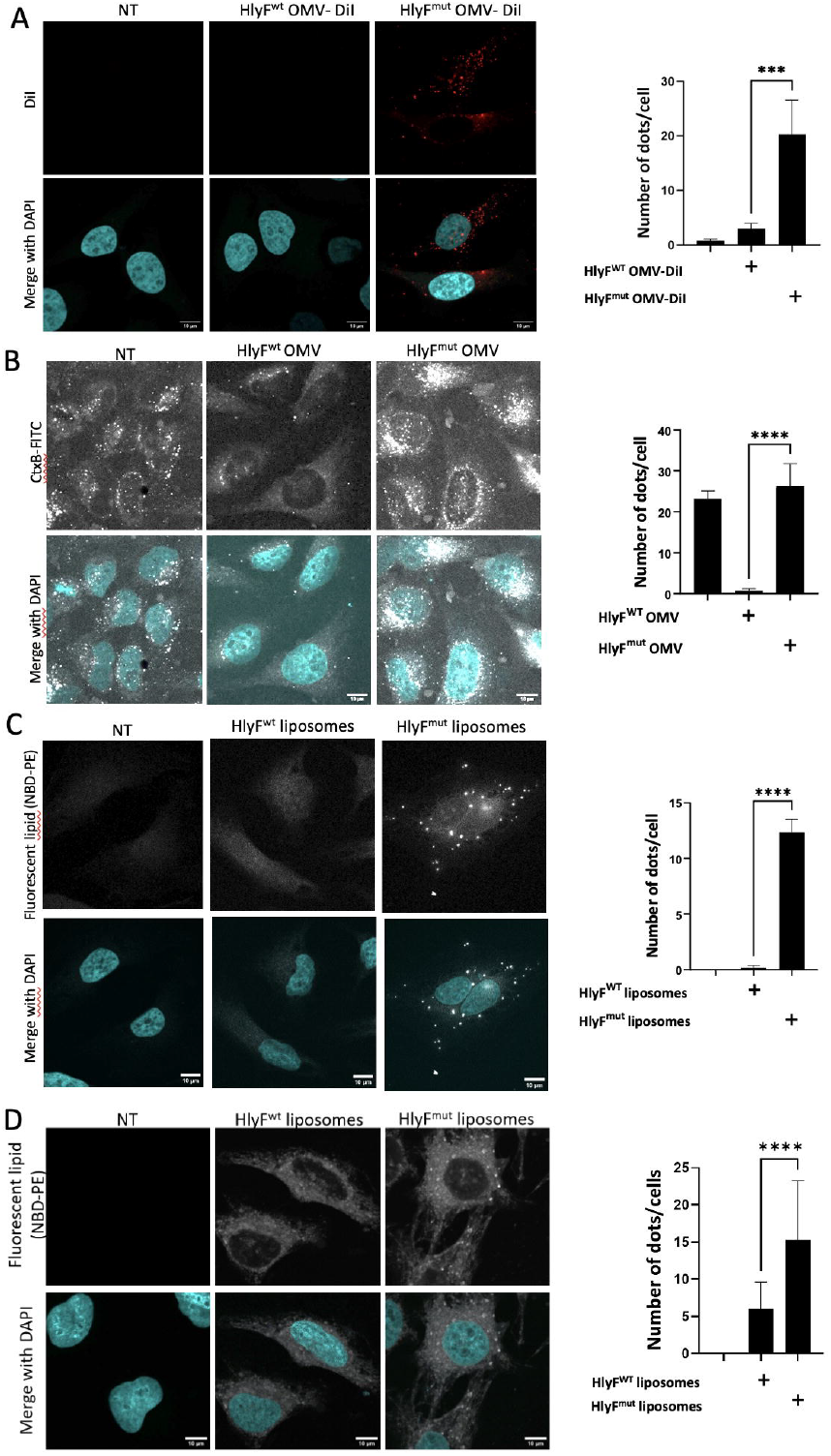
HlyF^wt^ OMVs are internalized in host cells by direct fusion to plasma membrane. (A) Confocal microscopy of HeLa cells treated for 2hr with 10µg/mL DiI-stained HlyF^wt^ or HlyF^mut^ OMVs. OMVs were visualized using DiI fluorescence (565nm, red). Images are representative of 3 independent experiments; (B) Confocal microscopy of HeLa cells treated for 1.5hr with 5µg/mL HlyF^wt^ or HlyF^mut^ OMVs. Lipid raft distribution was visualized by cholera toxin B subunit conjugated to FITC (CtxB-FITC). Images are representative of 5 independent experiments; (C) Confocal microscopy of HeLa cells treated for 6h with liposomes derived from either HlyF^wt^ or HlyF^mut^ expressing-strains (100µg of lipid/mL). Liposomes were visualized using NBD–phosphatidylethanolamine fluorescence (310nm). Images are representative of two independent experiments; (D) Confocal microscopy of HeLa cells treated for 4h with liposomes derived from either HlyF^wt^ or HlyF^mut^ OMVs (50µg of lipid/mL). Liposomes were visualized using NBD–phosphatidylethanolamine fluorescence (310nm). For each panel, nuclei were counterstained with DAPI; Scale bar, 10µm; at least 50 cells were analyzed per condition for quantification with the ImageJ software; the number of fluorescent particles (dots) into cells is indicated on the right panel for each experiment. The graphs show the mean and the standard deviation for each condition. *** p = 0,0003; **** p < 0.0001 Welch’s t-test.

### Lipid composition confers fusogenic capacity to HlyF OMVs

HlyF and CprA belong to a family of SDR enzymes that modify lipids. We previously showed that liposomes reconstituted from lipids of CprA-expressing OMVs were sufficient to inhibit autophagy, indicating that the active component resides within the lipid fraction (Goman et al. 2025). To assess whether the fusogenicity of OMVs from bacteria expressing these enzymes is also lipid-dependent, we generated fluorescent liposomes with lipids extracted from either the bacterial membranes or the purified OMVs, from BL21(DE3) strains expressing either HlyF^mut^ or HlyF^wt^. Upon incubation with HeLa cells, HlyF^wt^ derived liposomes produced diffuse fluorescence throughout the cell, whereas HlyF^mut^ derived liposomes displayed a punctate pattern, consistent with endosomal confinement. These observations were further corroborated by quantitative puncta analysis (Figure 5C and 5D). These results support a model in which SDR-mediated lipid remodeling by HlyF enables OMVs or liposomes to bypass endocytic uptake and directly fuse with the plasma membrane, whereas vesicles lacking SDR-modified lipids remain trapped in endosomal compartments.

## Discussion

We demonstrate that the HlyF/CprA enzymes drive the production of a distinct subpopulation of OMVs that deliver their cargo to host cells more rapidly and efficiently than OMVs derived from *hlyF*-negative strains. These findings provide strong evidence that the enzymatic activity of HlyF/CprA not only regulates OMV biogenesis but also critically determines how OMVs interact with eukaryotic membranes. This dual role likely underlies the specific functional properties of OMVs from HlyF-producing bacteria, including their capacity to interfere with autophagic flux.

While we could not exclude macropinocytosis as an additional uptake route, our results strongly support a model in which HlyF^wt^ OMVs bypass classical endocytosis and instead enter host cells via direct fusion with the plasma membrane. Importantly, this phenotype was recapitulated using reconstituted liposomes generated from lipids extracted from OMVs or bacterial membranes of HlyF^wt^ or HlyF^mut^ -expressing strains, demonstrating that vesicle lipid composition alone is sufficient to dictate this behavior. These observations identify HlyF-modified lipids as the primary drivers of this alternative entry mechanism, independent of vesicle-associated proteins or cargo. Fusion-mediated internalization provides a compelling mechanistic explanation for the rapid and extensive cytosolic delivery of OMV cargo observed with HlyF^wt^ vesicles. Unlike endocytosis, which requires vesicular trafficking and endosomal maturation prior to cargo release (Doherty and McMahon 2009; Kumari, Mg, and Mayor 2010), membrane fusion enables immediate transfer of OMV vesicular luminal contents into the host cell cytosol. This mechanism is consistent with observations in *P. aeruginosa* strains expressing CprA, the HlyF ortholog, where OMVs similarly enter host cells through a dynamin-independent, fusion-based pathway (Bomberger et al. 2009). These findings suggest that lipid-driven OMV fusion may represent a conserved strategy among HlyF/CprA-positive pathogens.

At the molecular level, both bacterial and eukaryotic membrane dynamics are governed by lipid composition, which determines key physical properties such as curvature, rigidity, packing, density and phase behavior, parameters that critically influence miscibility and fusogenic potential (Rietveld and Simons 1998). Recent work has identified HlyF/CprA as members of the SDR enzymes that modify lipids that become incorporated into OMV lipids (Goman et al. 2025). Although the precise lipid species modified by HlyF/CprA remain to be identified, such enzymatic remodeling would be expected to alter bilayer flexibility and reduce membrane stability, thereby lowering the energetic barrier to fusion with host membranes (Chernomordik and Kozlov 2008). By reprogramming OMV lipid composition, HlyF/CprA may thus selectively enhance vesicle compatibility with lipid raft microdomains, including cholesterol, consistent with the observed disruption of CtxB-positive domains.

These findings have important implications for bacterial virulence. Clinical *E. coli and P. aeruginosa* strains expressing HlyF/CprA produce OMVs that efficiently deliver a broad range of virulence factors, including Shiga toxin, cytolethal distending toxin and the CFTR inhibitory factor (Bielaszewska et al. 2017; Bomberger et al. 2009; Soysal et al. 2016). Direct fusion of OMVs with the plasma membrane would allow these effectors to bypass endosomal sorting pathways, enabling their immediate release into the host cytosol and direct access to intracellular targets. Moreover, the concurrent inhibition of autophagic clearance by HlyF/CprA OMVs (David et al. 2022; Goman et al. 2025) would further prolong the intracellular persistence and activity of these toxins. The combination of enhanced delivery and impaired degradation likely contributes to the heightened cytotoxicity, inflammation, and disease severity observed during infections with *hlyF/cprA*-positive strains (Chagneau et al. 2023; Goman et al. 2025; Murase et al. 2016).

The identification of *hlyF/cprA* homologs in diverse pathogens, including *Yersinia pestis* and *Ralstonia solanacearum* (Goman et al. 2025), underscores the evolutionary conservation of this strategy and suggests that lipid remodeling to modulate OMV–host membrane interactions may be a widespread, yet underappreciated, virulence mechanism among Gram-negative bacteria. These findings expand the conceptual framework of OMV biology by highlighting that vesicle function is governed not only by cargo composition but also by lipid-driven entry mechanisms.

In conclusion, our study identifies HlyF/CprA as key enzymatic regulators of OMV lipid composition that confer fusogenic properties and enable a novel, dynamin-independent mode of host cell entry. This fusion-based mechanism provides a rapid and efficient pathway for bacterial cargo delivery and likely contributes substantially to the virulence of HlyF/CprA-positive pathogens. By linking lipid enzymology to vesicle–host membrane interactions, our work reveals a fundamental aspect of EV biology with broad implications for infection biology, host–pathogen interactions, and vesicle-based therapeutic design. Future studies aimed at defining the specific lipid species generated by HlyF/CprA and their effects on membrane phase behavior and fusion energetics will be essential to fully elucidate this mechanism.

## Materials and methods

### Bacterial strains and growth conditions

BL21(DE3) expressing wild-type and mutated HlyF were obtained following transformation with respectively, plasmids pAGO-15 and pAGO-16 previously described in (Goman et al. 2025). BL21(DE3) strains were further transformed with pUC19 plasmid for TEM-1 ß-lactamase expression (Addgene).

ECC166 is an uropathogenic *E. coli* strain (UPEC) isolated from woman with pyelonephritis. ECC166 and ECC166 *ΔhlyF* strains described in (Chagneau et al. 2023) were further transformed with pUC19 plasmid. *P. aeruginosa PAK ΔcprA and PAK ΔcprA* complemented with CprA strains were previously described in (Goman et al. 2025). PAK strains were further transformed with pJN-TEM plasmid in which the *tem-1* gene from pUC19 was inserted into the EcoRI-XbaI sites of pJN105 plasmid (Addgene) under the control of AraBAD promoter. For OMV production, bacteria were grown at 37°C under vigorous shaking (240rpm) for 8h in M63 medium supplemented with casamino-acid and appropriate antibiotic and inoculated with 1/100 of overnight culture. For PAK strains, 1% rhamnose and 1% arabinose were added after 3h of culture for CprA and TEM-1 induction respectively.

### Purification of bacterial OMVs

After 8h of culture, the bacterial suspension was centrifuged at 6500g for 10min at 4°C. The supernatant was filtered (pore size 0.45μm; Merck-Millipore) to obtain bacterial-free supernatant. The filtered supernatant was ultrafiltered and concentrated using tangential concentrator and PES membrane cassette (pore size 100kDa; Sartorius) and then ultracentrifuged at 150,000g for 2h at 4°C. After removing the supernatant, pelleted OMV were resuspended in sterile PBS and stored at 4°C. To visualize the concentration of OMVs and the purity of the preparation, negative staining transmission electron microscopy (TEM) was performed according to standard procedures. The concentration of OMVs in the suspension correlates with protein concentration evaluated by BCA protein assay (Bio-Rad, RC DCTM Protein Assay kit II). When indicated, pronase treatment was performed by incubating OMV (125µg/mL) with pronase (2mg/mL) for 90min at 37°C. Following digestion, samples were treated with cOmplete protease inhibitor (Roche) and 1mM phenylmethylsulfonyl fluoride (PMSF), then ultracentrifuged and resuspended in PBS at initial volume.

### Particle measurements

Particle number and size distribution in samples were determined by multi-angle particle analysis using a Zetasizer Advance Series - Ultra (Red Label) (Malvern Technologies, Malvern, UK). Samples were diluted (1:100 & 1:1000) in particle-free PBS to an appropriate concentration according to the manufacturer’s recommendations. Measurements were performed in triplicate for each dilution.

### Eukaryotic cell culture

HeLa cells (ATCC, HeLa CCL2) were cultured in DMEM with Glutamax (Life Technology) supplemented with 10% fetal bovine serum (FBS) and non-essential amino-acids (Life Technology). One day before the experiment, HeLa cells were seeded in 96-wells plate at a density of 10 cells per well in 100µL of DMEM supplemented with 10% FBS. One week before treatment, bone marrow-derived macrophages (BMDM) were prepared from fresh bone marrow isolated from C57BL/6 mice at a density of 2 × 10 cells per well. Bone marrow cells were allowed to differentiate in macrophages by incubation for 7 days in DMEM supplemented with 10% FBS, 25ng/ml CSF1/M-CSF (Immunotools), 10mM HEPES, pH 7.2 and nonessential amino acids (Life Technology).

### CCF4-AM assays

Twenty µL of 6X loading solution with CCF4-AM (LiveBLAzer FRET-B/G loading kit; Life technologies) supplemented with 2.5mM of probenicid were added to 100µL of cell culture in p96-well plate seeded at day-1 and further incubated at room temperature for 2h. Cells were rinsed 3 times with 100µl of HBSS and incubated with indicated concentration of OMVs in OptiMEM medium with probenicid. When indicated, Dynasore (80µM), chloroquine (100µM) or bafilomycin (100nM) were added together with OMVs. Background control wells without CCF4, with and without inhibitors were performed. Each condition was done in triplicate. Plate reading was performed with Tecan Infinite 200 Pro (Tecan) at 37°C, every 15min during 8hr with excitation set at 410nm and detection of emission at 450nm (blue fluorescence) and 520nm (green fluorescence). CCF4-AM cleavage was calculated by the ratio of the net blue fluorescence (-background) divided by the net green fluorescence.

### Immunoblotting

Detection of OMPA and TEM-1 ß-lactamase were performed with 2µg of purified OMVs preparation per lane, separated by SDS-PAGE (Invitrogen) and transferred to PVDF membranes (Invitrogen). The membranes were blocked in 5% non-fat dry milk in TBS 0.1% Tween® 20 and probed with rabbit anti *E. coli* OMPA antibody (Antibody Research Corporation) and mouse anti-beta lactamase antibody [8A5.A10] (Abcam). After incubation with goat anti-rabbit and goat anti-mouse HRP-conjugated secondary antibodies (Bethyl Laboratories), proteins were visualized using chemiluminescent peroxidase substrate detection reagents (Sigma), and acquired by the ChemiDoc XRS+ System (Bio-Rad). Protein quantification was performed by densitometric analyses of immunoblots using the ImageJ software. Of note, OMV samples were not heated after Laemli buffer addition to preserve OmpA as a single form (see Figure S4).

### Nitrocefin hydrolysis assay

Ten µL of OMVs at 50ng/mL were transferred into a 96-well microplate on ice with 40µL of PBS and 50µL of nitrocefin solution (Euromedex) at 10mg/mL. Each condition was done in triplicate.

Absorbance at 486nm was measured immediately using Infinite 200 Pro plate reader (Tecan) and subsequently every 5min for 2hr.

### Lipid extraction

For lipid extraction, we used a Folch-like extraction method (Folch, Lees, and Sloane Stanley 1957). Briefly, bacteria were grown under the same conditions as for OMV production (see OMV production above). After 8hr of culture at 37°C with agitation at 240rpm, the cultures were centrifuged at 6,000 × g for 20min to pellet the bacteria and remove the supernatant. The bacterial pellet or purified OMV (see purification of bacterial OMV section above) was then resuspended in a 2:1 CHCl_3_/MeOH solution and agitated for 1hr at 40rpm in a rotary shaker. Samples were then centrifugated for 10min à 1,000rpm. The supernatants were then washed with water and agitated for 15min at 40rpm in a rotary shaker. Samples were then centrifugated for 10min at 1,000rpm and the organic phases were recovered and evaporated under N_2_. After weighing, dry lipids pellets were resuspended in DMEM without phenol red (21063 Gibco). Lipid concentrations were quantified with a sulfo-phospho-vanillin reaction as described in Izard et al. *(Izard and Limberger 2003)* with Triolein as reference (44896-U Sigma Aldrich).

### Liposomes preparation

Liposomes were generated from mix of lipids derived from HlyF-expressing bacteria or purified OMVs (extracted as described above) and a commercial fluorescent lipid, namely 1-palmitoyl-2 -(Schwechheimer and Kuehn 2015)-sn-glycero-3-phosphoethanolamine (NBD-PE, 810154P-1MG Avanti), adapted from Kehl et al. *(Kehl et al. 2020)*. Before extrusion, lipidic mixes were prepared with 1mg of lipids from HlyF^wt^ - or from HlyF^mut^ -expressing bacteria and 0.6mg of 16:0-12:0 NBD PE, 0.3mg of 1-palmitoyl-2-oleoyl-sn-glycero-3-phosphoglycerol and 0.1mg of cardiolipin resuspended in chloroform/methanol (2/1, v/v). Lipidic mixes were then evaporated under N_2._ After weighing, dry lipids pellets were resuspended in DMEM without phenol red (Gibco). Liposomes were then generated by extrusion, according to the manufacturer’s instruction about the Avanti Mini Extruder Extrusion Technique (610000-1EA Avanti Polar Lipids Inc.). Lipids mixes were extruded using 200nm (610006-1EA Avanti) then 100nm membranes (610005-1EA Avanti) and stored at 4°C. Images were acquired using a Zeiss LSM 710 confocal microscope. Images were subsequently processed using the ImageJ software package.

### DiI labelling

OMVs samples (5mg/mL) were labelled with 1% (v:v) DiI (Invitrogen, V22885) for 30min at 37 °C and then immediately directly added to the medium of HeLa cells culture without any additional step.

### Fluorescence microscopy

HeLa cells were seeded at a density of 5 × 10^5^ cells per well in 1mL of DMEM supplemented with 10% FBS into 12-well plates with slides in the bottom of the wells the day before the experiment. The day of the experiment, cells were treated with OMVs or liposomes at the indicated concentration and time. When indicated, CtxB-FITC (Sigma, C1655) was added directly in the medium of cell culture at a final concentration of 1µg/mL at the same time as OMV. At the end of the experiment, the cells were fixed with 4% PFA for 12min at RT. After being washed 3 times with PBS the slides were mounted with Fluoroshield (sigma F6182). Images were acquired using a Zeiss LSM 710 confocal microscope. Images were subsequently processed using the ImageJ software package.

### Statistical analyses

All graphical figures and statistical analyses were performed with the GraphPad Prism 11 software (GraphPad Software, San Diego, CA).

## Abbreviations

AUC: area under the curve CtxB: cholera toxin B
*E. coli*: *Escherichia coli*
EV(s): Extracellular vesicle(s)
FRET: fluorescence resonance energy transfer HlyF: hemolysin F
LPS: lipopolysaccharide
MADLS: multi angle dynamic light scattering
mut: mutated
M/PAMPs: microbial/pathogen-associated molecular patterns
NTA: nanoparticle tracking analysis
OMP(s): outer membrane protein(s)
OMV(s): outer membrane vesicle(s)
*P. aeruginosa*: *Pseudomonas aeruginosa*
SDR: short chain dehydrogenase/reductase
TEM: transmission electron microscopy
UPEC: uropathogenic *E. coli*
wt: wild-type

## References

Arab, T., E. R. Mallick, Y. Huang, L. Dong, Z. Liao, Z. Zhao, O. Gololobova, B. Smith, N. J. Haughey, K. J. Pienta, B. S. Slusher, P. M. Tarwater, J. P. Tosar, A. M. Zivkovic, W. N. Vreeland, M. E. Paulaitis, and K. W. Witwer. 2021. “Characterization of extracellular vesicles and synthetic nanoparticles with four orthogonal single-particle analysis platforms.” J Extracell Vesicles 10 (6): e12079.

Bachurski, D., M. Schuldner, P. H. Nguyen, A. Malz, K. S. Reiners, P. C. Grenzi, F. Babatz, A. C. Schauss, H. P. Hansen, M. Hallek, and E. Pogge von Strandmann. 2019. “Extracellular vesicle measurements with nanoparticle tracking analysis - An accuracy and repeatability comparison between NanoSight NS300 and ZetaView.” J Extracell Vesicles 8 (1): 1596016.

Bielaszewska, M., C. Ruter, A. Bauwens, L. Greune, K. A. Jarosch, D. Steil, W. Zhang, X. He, R. Lloubes, A. Fruth, K. S. Kim, M. A. Schmidt, U. Dobrindt, A. Mellmann, and H. Karch. 2017. “Host cell interactions of outer membrane vesicle-associated virulence factors of enterohemorrhagic Escherichia coli O157: Intracellular delivery, trafficking and mechanisms of cell injury.” PLoS Pathog 13 (2): e1006159.

Bomberger, J. M., D. P. Maceachran, B. A. Coutermarsh, S. Ye, G. A. O’Toole, and B. A. Stanton. 2009. “Long-distance delivery of bacterial virulence factors by Pseudomonas aeruginosa outer membrane vesicles.” PLoS Pathog 5 (4): e1000382.

Canas, M. A., R. Gimenez, M. J. Fabrega, L. Toloza, L. Baldoma, and J. Badia. 2016. “Outer Membrane Vesicles from the Probiotic Escherichia coli Nissle 1917 and the Commensal ECOR12 Enter Intestinal Epithelial Cells via Clathrin-Dependent Endocytosis and Elicit Differential Effects on DNA Damage.” PLoS One 11 (8): e0160374.

Chagneau, C. V., D. Payros, A. Goman, C. Goursat, L. David, M. Okuno, P. J. Bordignon, C. Seguy, C. Massip, P. Branchu, Y. Ogura, J. P. Nougayrede, M. Marenda, and E. Oswald. 2023. “HlyF, an underestimated virulence factor of uropathogenic Escherichia coli.” Clin Microbiol Infect 29 (11): 1449 e1–1449 e9.

Chernomordik, L. V., and M. M. Kozlov. 2008. “Mechanics of membrane fusion.” Nat Struct Mol Biol 15 (7): 675–83.

David, L., F. Taieb, M. Penary, P. J. Bordignon, R. Planes, S. Bagayoko, V. Duplan-Eche, E. Meunier, and E. Oswald. 2022. “Outer membrane vesicles produced by pathogenic strains of Escherichia coli block autophagic flux and exacerbate inflammasome activation.” Autophagy 18 (12): 2913–2925.

Doherty, G. J., and H. T. McMahon. 2009. “Mechanisms of endocytosis.” Annu Rev Biochem 78: 857–902.

Eren, E., R. Planes, S. Bagayoko, P. J. Bordignon, K. Chaoui, A. Hessel, K. Santoni, M. Pinilla, B. Lagrange, O. Burlet-Schiltz, J. C. Howard, T. Henry, M. Yamamoto, and E. Meunier. 2020. “Irgm2 and Gate-16 cooperatively dampen Gram-negative bacteria-induced caspase-11 response.” EMBO Rep 21 (11): e50829.

Folch, J., M. Lees, and G. H. Sloane Stanley. 1957. “A simple method for the isolation and purification of total lipides from animal tissues.” J Biol Chem 226 (1): 497–509.

Goman, A., B. Ize, K. Jeannot, C. Pin, D. Payros, C. Goursat, L. Ravon-Katossky, K. Murase, C. V. Chagneau, H. Revillet, F. Taieb, S. Bleves, L. David, E. Meunier, P. Branchu, and E. Oswald. 2025. “Uncovering a new family of conserved virulence factors that promote the production of host-damaging outer membrane vesicles in gram-negative bacteria.” J Extracell Vesicles 14 (1): e270032.

Harder, T., P. Scheiffele, P. Verkade, and K. Simons. 1998. “Lipid domain structure of the plasma membrane revealed by patching of membrane components.” J Cell Biol 141 (4): 929–42.

Izard, J., and R. J. Limberger. 2003. “Rapid screening method for quantitation of bacterial cell lipids from whole cells.” J Microbiol Methods 55 (2): 411–8.

Jan, A. T. 2017. “Outer Membrane Vesicles (OMVs) of Gram-negative Bacteria: A Perspective Update.” Front Microbiol 8: 1053.

Juodeikis, R., and S. R. Carding. 2022. “Outer Membrane Vesicles: Biogenesis, Functions, and Issues.” Microbiol Mol Biol Rev 86 (4): e0003222.

Kaderabkova, N., M. Bharathwaj, R. C. D. Furniss, D. Gonzalez, T. Palmer, and D. A. I. Mavridou. 2022. “The biogenesis of beta-lactamase enzymes.” Microbiology (Reading) 168 (8).

Kehl, A., R. Kuhn, J. Detzner, D. Steil, J. Muthing, H. Karch, and A. Mellmann. 2020. “Modeling Native EHEC Outer Membrane Vesicles by Creating Synthetic Surrogates.” Microorganisms 8 (5).

Kesty, N. C., K. M. Mason, M. Reedy, S. E. Miller, and M. J. Kuehn. 2004. “Enterotoxigenic Escherichia coli vesicles target toxin delivery into mammalian cells.” EMBO J 23 (23): 4538–49.

Krsek, D., D. A. Yara, H. Hrbackova, O. Daniel, A. Mancikova, S. Schuller, and M. Bielaszewska. 2023. “Translocation of outer membrane vesicles from enterohemorrhagic Escherichia coli O157 across the intestinal epithelial barrier.” Front Microbiol 14: 1198945.

Kumari, S., S. Mg, and S. Mayor. 2010. “Endocytosis unplugged: multiple ways to enter the cell.” Cell Res 20 (3): 256–75.

Kunsmann, L., C. Ruter, A. Bauwens, L. Greune, M. Gluder, B. Kemper, A. Fruth, S. N. Wai, X. He, R. Lloubes, M. A. Schmidt, U. Dobrindt, A. Mellmann, H. Karch, and M. Bielaszewska. 2015. “Virulence from vesicles: Novel mechanisms of host cell injury by Escherichia coli O104:H4 outbreak strain.” Sci Rep 5: 13252.

Lapaquette, P., J. Guzzo, L. Bretillon, and M. A. Bringer. 2015. “Cellular and Molecular Connections between Autophagy and Inflammation.” Mediators Inflamm 2015: 398483.

Macia, E., M. Ehrlich, R. Massol, E. Boucrot, C. Brunner, and T. Kirchhausen. 2006. “Dynasore, a cell-permeable inhibitor of dynamin.” Dev Cell 10 (6): 839–50.

Marks, B., M. H. Stowell, Y. Vallis, I. G. Mills, A. Gibson, C. R. Hopkins, and H. T. McMahon. 2001. “GTPase activity of dynamin and resulting conformation change are essential for endocytosis.” Nature 410 (6825): 231–5.

Murase, K., P. Martin, G. Porcheron, S. Houle, E. Helloin, M. Penary, J. P. Nougayrede, C. M. Dozois, T. Hayashi, and E. Oswald. 2016. “HlyF Produced by Extraintestinal Pathogenic Escherichia coli Is a Virulence Factor That Regulates Outer Membrane Vesicle Biogenesis.” J Infect Dis 213 (5): 856–65.

Naiim, M., A. Boualem, C. Ferre, M. Jabloun, A. Jalocha, and P. Ravier. 2015. “Multiangle dynamic light scattering for the improvement of multimodal particle size distribution measurements.” Soft Matter 11 (1): 28–32.

Normak, K., M. Papp, M. Ullmann, C. Paganini, M. Manno, A. Bongiovanni, P. Bergese, and P. Arosio. 2023. “Multiparametric Orthogonal Characterization of Extracellular Vesicles by Liquid Chromatography Combined with In-Line Light Scattering and Fluorescence Detection.” Anal Chem 95 (33): 12443–12451.

O’Donoghue, E. J., and A. M. Krachler. 2016. “Mechanisms of outer membrane vesicle entry into host cells.” Cell Microbiol.

Park, K. S., K. H. Choi, Y. S. Kim, B. S. Hong, O. Y. Kim, J. H. Kim, C. M. Yoon, G. Y. Koh, Y. K. Kim, and Y. S. Gho. 2010. “Outer membrane vesicles derived from Escherichia coli induce systemic inflammatory response syndrome.” PLoS One 5 (6): e11334.

Pathirana, R. D., and M. Kaparakis-Liaskos. 2016. “Bacterial membrane vesicles: Biogenesis, immune regulation and pathogenesis.” Cell Microbiol.

Pin, C., L. David, and E. Oswald. 2023. “Modulation of Autophagy and Cell Death by Bacterial Outer-Membrane Vesicles.” Toxins (Basel) 15 (8).

Rietveld, A., and K. Simons. 1998. “The differential miscibility of lipids as the basis for the formation of functional membrane rafts.” Biochim Biophys Acta 1376 (3): 467–79.

Sabatke, B., I. V. Rossi, A. Sana, L. B. Bonato, and M. I. Ramirez. 2024. “Extracellular vesicles biogenesis and uptake concepts: A comprehensive guide to studying host-pathogen communication.” Mol Microbiol 122 (5): 613–629.

Santos, J. C., M. S. Dick, B. Lagrange, D. Degrandi, K. Pfeffer, M. Yamamoto, E. Meunier, P. Pelczar, T. Henry, and P. Broz. 2018. “LPS targets host guanylate-binding proteins to the bacterial outer membrane for non-canonical inflammasome activation.” EMBO J 37 (6).

Schwechheimer, C., and M. J. Kuehn. 2015. “Outer-membrane vesicles from Gram-negative bacteria: biogenesis and functions.” Nat Rev Microbiol 13 (10): 605–19.

Soysal, N., P. Mariani-Kurkdjian, Y. Smail, S. Liguori, M. Gouali, E. Loukiadis, P. Fach, M. Bruyand, J. Blanco, P. Bidet, and S. Bonacorsi. 2016. “Enterohemorrhagic Escherichia coli Hybrid Pathotype O80:H2 as a New Therapeutic Challenge.” Emerg Infect Dis 22 (9): 1604–12.

Stamenkovic, M., K. Janjetovic, V. Paunovic, D. Ciric, T. Kravic-Stevovic, and V. Trajkovic. 2019. “Comparative analysis of cell death mechanisms induced by lysosomal autophagy inhibitors.” Eur J Pharmacol 859: 172540.

Thapa, H. B., P. Kohl, F. G. Zingl, D. Fleischhacker, H. Wolinski, T. A. Kufer, and S. Schild. 2023. “Characterization of the Inflammatory Response Evoked by Bacterial Membrane Vesicles in Intestinal Cells Reveals an RIPK2-Dependent Activation by Enterotoxigenic Escherichia coli Vesicles.” Microbiol Spectr 11 (4): e0111523.

Vanaja, S. K., A. J. Russo, B. Behl, I. Banerjee, M. Yankova, S. D. Deshmukh, and V. A. K. Rathinam. 2016. “Bacterial Outer Membrane Vesicles Mediate Cytosolic Localization of LPS and Caspase-11 Activation.” Cell 165 (5): 1106–1119.

