## Supplemental files for "HlyF/CprA modulate bacterial extracellular vesicle properties and mediate efficient host cell cargo delivery"

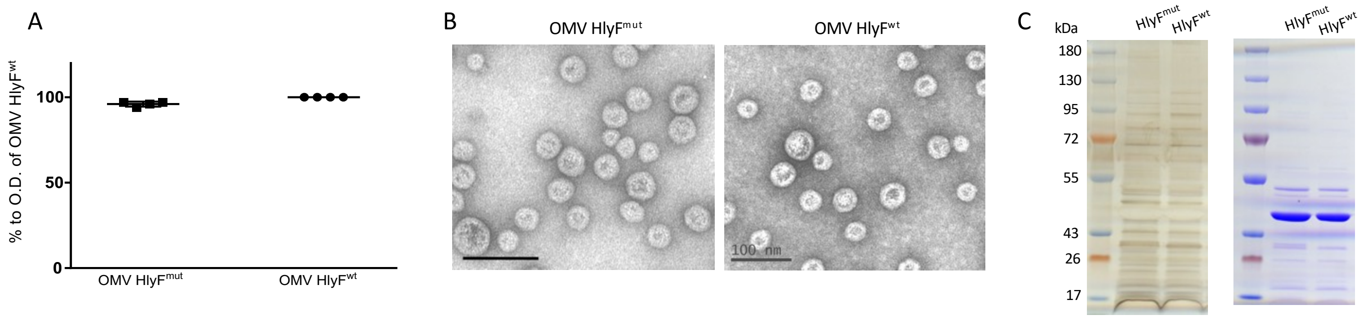


**Supplementary figure 1. Characterization of OMV produced by BL21(DE3) expressing wild-type or mutated HlyF.**

**(A)** Quantification of ß-lactamase activity after 2hr incubation of HlyF$^{mut}$ and HlyF$^{wt}$ OMVs with nitrocefin with OMVs samples normalized by western blot, from 4 independent experiments

**(B)** Negative staining transmission electronic microscopy of purified HlyF$^{mut}$ and HlyF$^{wt}$ OMVs (scale bar 100nm)

**(C)** Silver (left) and coomassie blue (right) staining of HlyF$^{mut}$ and HlyF$^{wt}$ OMV proteins runed on SDS-PAGE

A

| | DIAMETER (nm) | | | | | | | | | | | | | |
|---|---|---|---|---|---|---|---|---|---|---|---|---|---|---|
| | pop1 | SD | pop2 | SD | pop3 | SD | pop4 | SD | | | | | | |
| OMV WT 29/4 | 32.45 | 1.31 | 104.01 | 10.46 | 0.00 | 0.00 | 0.00 | 0.00 | | | | | | |
| OMV mut 29/4 | 27.15 | 2.48 | 96.23 | 9.77 | 0.00 | 0.00 | 0.00 | 0.00 | | | | | | |
| OMV WT 29/4 + O | 34.05 | 2.29 | 102.80 | 3.60 | 0.00 | 0.00 | 0.00 | 0.00 | | | | | | |
| OMV mut 29/4 + O | 28.20 | 0.00 | 105.37 | 6.67 | 0.00 | 0.00 | 0.00 | 0.00 | | | | | | |
| OMV WT 6/5 | 27.26 | 1.41 | 80.50 | | 185.33 | 22.24 | 435.78 | 11.59 | | | | | | |
| OMV mut 6/5 | 22.89 | 1.13 | 69.66 | 8.36 | 0.00 | 0.00 | 0.00 | 0.00 | | | | | | |
| OMV WT 2025 | 28.97 | 1.70 | 153.00 | 15.10 | | | 447.80 | 20.90 | | | | | | |
| OMV mut 2025 | 24.27 | 2.18 | 92.61 | 10.75 | | | 452.13 | 30.40 | | | | | | |
| mean OMV WT | <u>29.56</u> | 2.64 | 112.50 | 30.20 | | | | | | | | | | |
| mean OMV mut | <u>24.77</u> | 2.17 | 86.17 | 11.77 | | | | | | | | | | |
| mean (all OMV) | <u>28.16</u> | 3.75 | | | | | | | | | | | | |
| | CONCENTRATION (particle/mL) | | | | | | | | protein concentration (mg/mL) | NORMALIZED CONCENTRATION (particle/mg of protein) | | | | |
| | pop1 | SD | pop2 | SD | pop3 | SD | pop4 | SD | | pop1 | SD | pop2 | SD | % pop2/pop1 |
| OMV WT 29/4 | 4.82E+15 | 6.78E+14 | 3.36E+12 | 1.68E+12 | 0 | 0 | 0 | 0 | 2.1 | 2.30E+15 | 3.23E+14 | 1.60E+12 | 8.00E+11 | 0.07 |
| OMV mut 29/4 | 4.352E+15 | 1.384E+15 | 1.89E+13 | 1.33E+13 | 0 | 0 | 0 | 0 | 1.8 | 2.42E+15 | 7.69E+14 | 1.05E+13 | 7.39E+12 | 0.43 |
| OMV WT 29/4 + O | 5.63E+15 | 2.41E+15 | 2.33E+12 | 6.25E+11 | 0 | 0 | 0 | 0 | 1.9 | 2.96E+15 | 1.27E+15 | 1.23E+12 | 3.29E+11 | 0.04 |
| OMV mut 29/4 + O | 2.26E+15 | 3.40E+14 | 8.50E+12 | 2.49E+12 | 0 | 0 | 0 | 0 | 1.6 | 1.41E+15 | 2.13E+14 | 5.31E+12 | 1.56E+12 | 0.38 |
| OMV WT 6/5 | 3.31E+15 | 1.03E+15 | 1.14E+12 | 0 | 1.33E+11 | 5.88E+10 | 3.27E+10 | 1.90E+10 | 1.3 | 2.55E+15 | 7.92E+14 | 8.77E+11 | 0.00E+00 | 0.04 |
| OMV mut 6/5 | 8.81E+15 | 8.30E+14 | 6.52E+13 | 5.51E+13 | 0 | 0 | 0 | 0 | 1.4 | 6.29E+15 | 5.93E+14 | 4.66E+13 | 3.94E+13 | 0.74 |
| OMV WT 2025 | 7.59E+15 | 2.67E+15 | 1.48E+12 | 1.16E+12 | | | | | 2.5 | 3.04E+15 | 1.07E+15 | 5.92E+11 | 4.64E+11 | 0.02 |
| OMV mut 2025 | 4.76E+15 | 2.43E+15 | 1.90E+13 | 1.45E+13 | | | | | 2.5 | 1.90E+15 | 9.72E+14 | 7.60E+12 | 5.80E+12 | 0.40 |
| mean (OMV WT) | | | | | | | | | | <u>2.71E+15</u> | 3.5E+14 | 1.07E+12 | 4.16E+11 | 0.04 |
| mean (OMV mut) | | | | | | | | | | <u>3.01E+15</u> | 2.23E+15 | 1.75E+13 | 1.95E+13 | 0.49 |
| mean (all OMV) | | | | | | | | | | <u>2.86E+15</u> | 3.6E+14 | 9.28E+12 | 1.55E+13 | <u>0.27</u> |

B

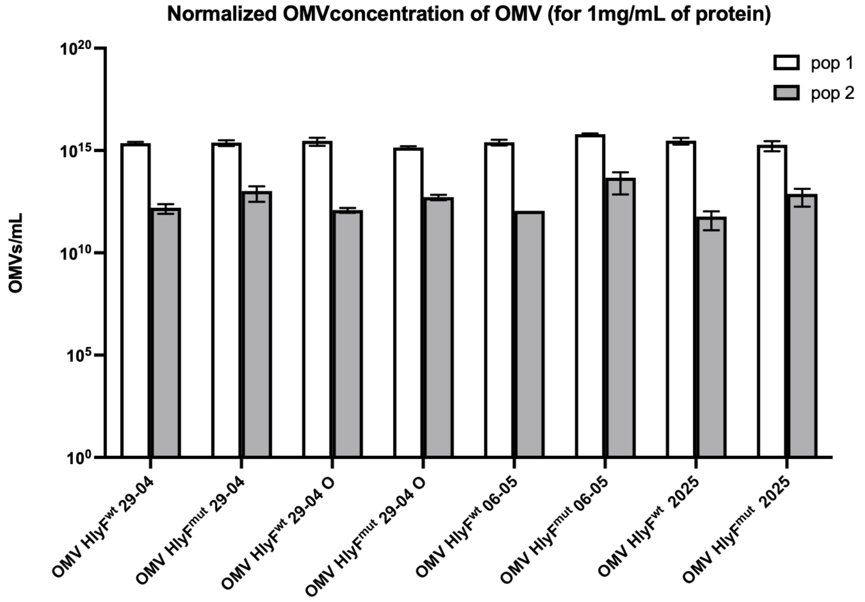


C

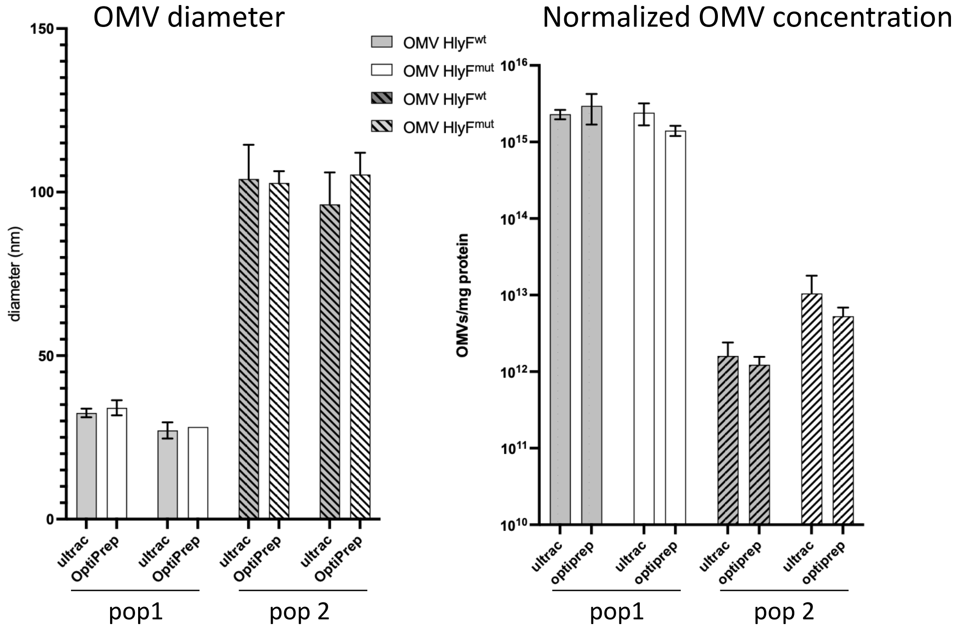


**Supplementary Figure 2. MADLS analysis of OMV size distribution and concentration**.

**(A)** Table summarizing MADLS measurements from three independent OMV purification batches (29-04, 06-05, and 2025) for HlyF$^{wt}$ OMVs (OMV WT) and HlyF$^{mut}$ OMVs (OMV mut) and after further purification by OptiPrep density-gradient separation for batch 29-04 (+O). Four populations (pop) were defined based on particle diameter: pop1, <50 nm; pop2, 50–175 nm; pop3, 175–400 nm; pop4, >400 nm. Vesicle sizes for each sample and mean values for HLyF$^{wt}$ OMVs and HlyF$^{mut}$ OMVs samples combined are reported as mean ± SD (top section). Vesicle concentrations were normalized to the protein concentration for each sample and the percentage of pop2 relative to pop1 was indicated; mean normalized concentrations and mean percentages were subsequently derived for HLyF$^{wt}$ OMVs, HlyF$^{mut}$ OMVs, and all OMVs samples combined (bottom section). Underlined values correspond to those reported in the main text.

**(B)** Column representation of the values presented in Table 1, showing side-by-side the concentrations of OMVs with diameters <50 nm (pop1) and >50 nm (pop2) for each purification batch. Data are presented as mean ± SD.

**(C)** MADLS analysis of HlyFwt (grey bars) and HlyFmut (white bars) OMVs, pop1 and pop 2 (hatched bars) showing particle diameter (left panel) and normalized particle concentration (OMV-to-protein ratio; right panel) after ultracentrifugation (ultrac) and following additional purification by density-gradient separation (OptiPrep). Data are presented as mean ± SD.

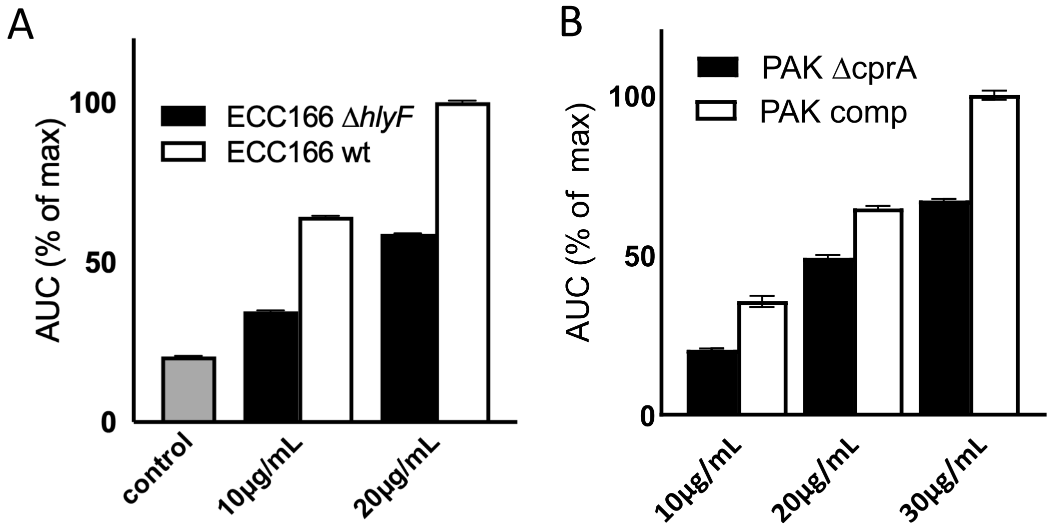


**Supplementary figure 3. OMVs from clinical pathogenic strains expressing HlyF/CprA promote cargo accumulation in host cells.**

AUC values representing total delivery of OMV-associated TEM-1 β-lactamase were calculated from time-course experiments as in figure 1D (technical triplicate experiment).

**(A)** HeLa cells treated or not (control, grey bar) with 10 or 20µg/mL OMVs from ECC166 Δ*hlyF* (black bars) or *wt* (white bars) strains.

**(B)** HeLa cells treated with 10, 20, or 30µg/mL OMVs from *P. aeruginosa* PAK Δ*cprA* (black bars) or PAK Δ*cprA* complemented with *cprA* (PAK comp) (white bars) strains.

AUC are represented as mean ± standard deviation (SD).

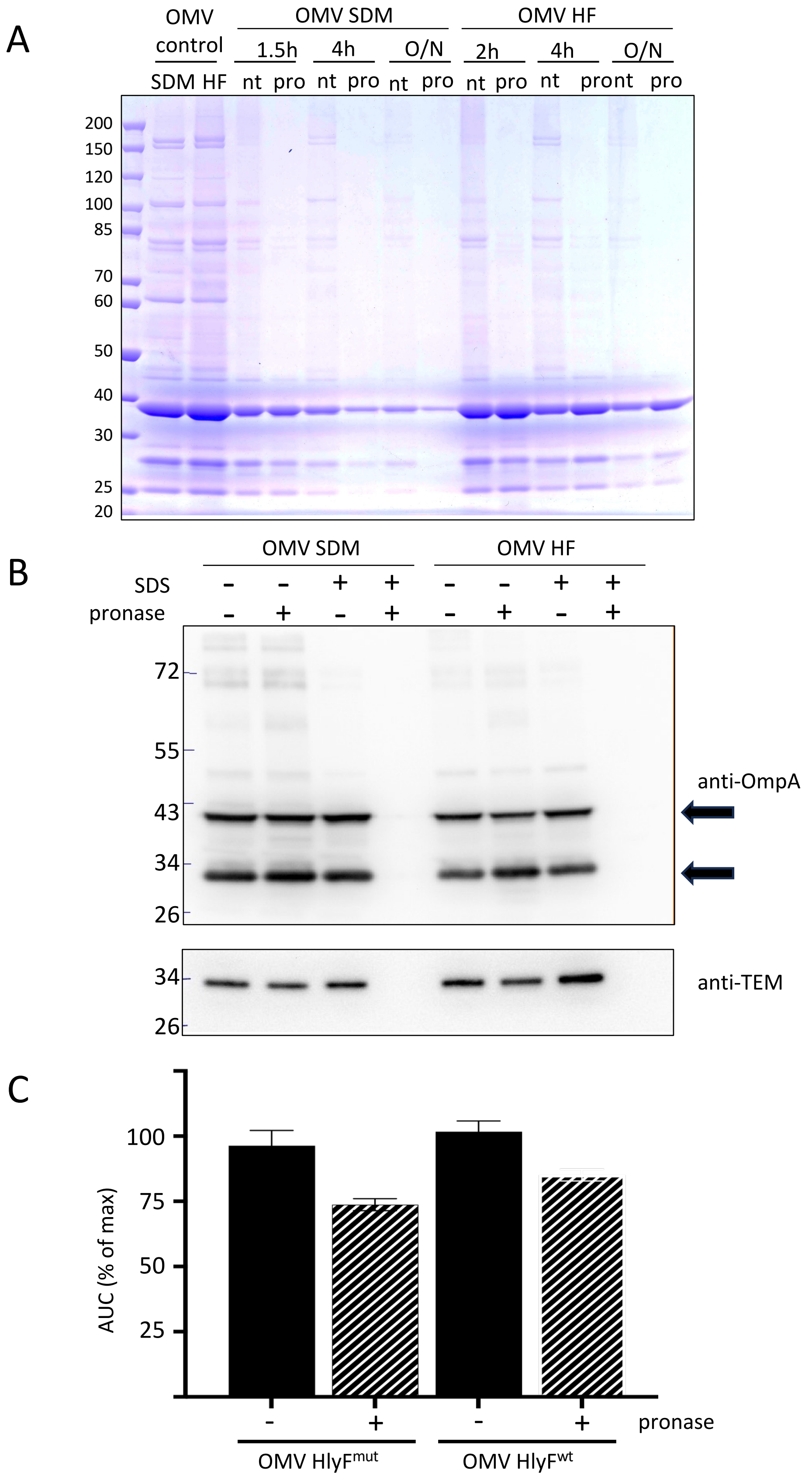


**Supplementary figure 4. OMV protect ß-lactamase from pronase proteolytic activity.**

**(A)**: OMVs (10µg) were incubated with (+) or without (-) with pronase for 1.5hr, 4hr or 18hr (O/N) then treated with PMSF and cOmplete protease inhibitor (Roche) and ultracentrifuged before loading on SDS-PAGE and protein staining; OMV control corresponds to OMV before treatment.

**(B)**: OmpA and TEM-1 ß-lactamase western blot detection of OMVs (1µg) incubated with (+) or without (-) pronase and SDS for 1.5hr then treated with PMSF and cOmplete protease inhibitor; Arrows indicated the two forms of OmpA after sample heating as described in Manning PA, Reeves P. Outer membrane of Escherichia coli K-12: differentiation of proteins 3A and 3B on acrylamide gels and further characterization of con (tolG) mutants. J Bacteriol. 1976 Sep;127(3):1070-9. Of note, samples were not heated for western blot figure 1B and OmpA quantification.

**(C)**: AUC from CCF4 assay with HeLa cells incubated with HlyF$^{mut}$ OMVs (25µg/mL) or HlyF$^{wt}$ (10µg/mL) incubated 90min at 37°C with (+) or whitout (-) pronase (2mg/mL), then treated with PMSF and cOmplete protease inhibitor and ultracentrifuged. AUC are presented as mean ± standard deviation (SD).

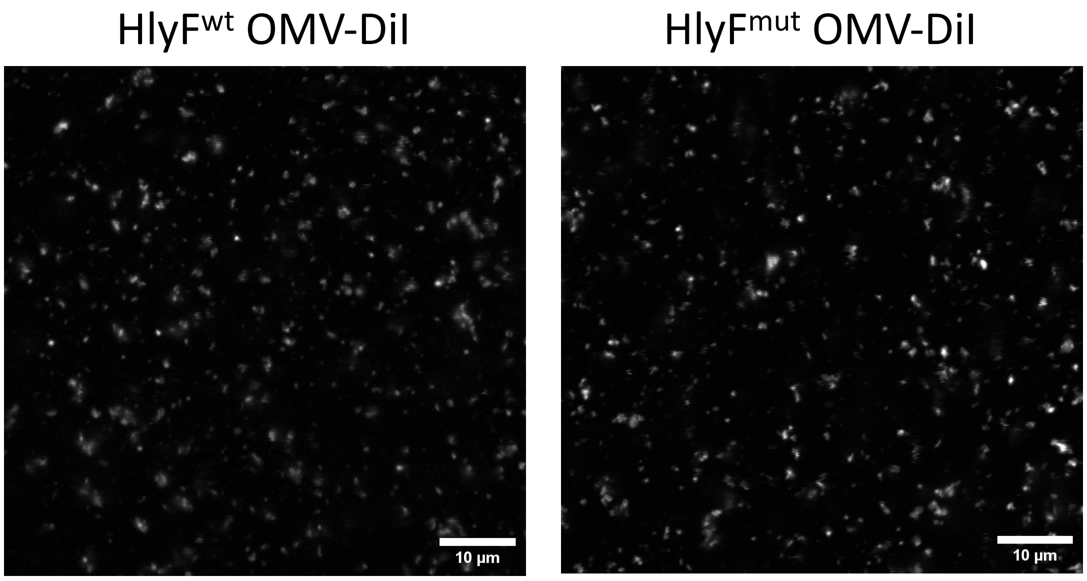


**Supplementary figure 5. DiI staining of OMVs.**
Confocal microscopy images of BL21(DE3) HlyF$^{wt}$ or HlyF$^{mut}$ OMVs stained with DiI. Puncta represent OMV-associated diffraction fluorescent signal Scale bar, 10µm.
